# Dysregulated splenic glucocorticoid sensitivity in aging and an α-synuclein transgenic mouse model of Parkinson’s disease

**DOI:** 10.64898/2026.08.27.745197

**Authors:** Daniel Rombach, Verena Bopp, Dominik Langgartner, Veselin Grozdanov, Jan Kassubek, Chadi Touma, Stefan O. Reber, Karin M Danzer

## Abstract

**Introduction:** Parkinson’s disease (PD) and aging both disrupt hypothalamic– pituitary–adrenal (HPA) axis function and peripheral immune homeostasis. Whether aging or α-synuclein (α-syn) pathology alters glucocorticoid (GC) sensitivity of peripheral immune cells has not been investigated.

**Methods:** Using an *ex vivo* GC sensitivity assay, we assessed the responsiveness of isolated and lipopolysaccharide (LPS)-stimulated splenocytes to the anti-inflammatory effects of increasing doses of corticosterone (CORT) in a wild-type (WT) aging cohort and in a PD α-syn transgenic mouse model and respective age-matched controls.

**Results:** Compared with splenocytes from 6-month-old WT mice, splenocytes from 20-month-old WT mice were less sensitive to 0.1 and 0.5 µM CORT. Isolated splenocytes from PD vs. control mice were less sensitive to 0.05, 0.1, and 0.5 µM CORT specifically at 16 months of age, but not at 6 or 20 months of age. As peripheral immune phenotyping revealed neither differences in HPA axis-related parameters nor in splenic GC receptor expression between PD and age-matched control mice at 6, 16, and 20 months, splenic GC resistance in PD mice at 16 months of age seems to be mediated by downstream GR signaling dysfunction.

**Conclusion:** Together, our results support the hypothesis that α-syn pathology accelerates an aging-associated decline in the peripheral sensitivity to anti-inflammatory GCs and may thereby sustain systemic and neuroinflammatory processes in PD.

## INTRODUCTION

Parkinson’s disease (PD) is one of the fastest-growing neurodegenerative disorders worldwide. Its prevalence doubles with each decade after the age of 65, and approximately 10 million people are currently affected worldwide [1,2]. The pathological hallmarks comprise the selective loss of dopaminergic neurons in an ascending propagation pattern, including in the substantia nigra pars compacta, and the accumulation of misfolded α-synuclein (α-syn) in Lewy bodies and Lewy neurites [3,4]. The resulting disruption of basal ganglia circuitry underlies the classical motor symptoms of PD, including bradykinesia, resting tremor, rigidity, and postural instability. However, PD is now recognized as a multisystem disorder in which non-motor symptoms, such as cognitive impairment, sleep disturbances, autonomic dysfunction, and neuropsychiatric manifestations, contribute substantially to disease burden and often precede motor symptom onset [1,5].

Among these non-motor neuropsychiatric features, depression affects approximately 40% of PD patients and represents one of the most prevalent and clinically relevant symptoms. Depression frequently emerges during the prodromal phase of PD, which suggests that it reflects underlying disease mechanisms rather than merely constituting a reactive consequence of neurodegeneration [6]. Increasing evidence points to dysregulation of the hypothalamic–pituitary–adrenal (HPA) axis as a central link between PD and depression [7]. Elevated cortisol levels and impaired negative feedback regulation have been consistently reported in PD patients, indicating disrupted stress-axis homeostasis [6,8]. In parallel, α-syn pathology extends beyond classical motor regions and affects several HPA axis-associated structures, including the hippocampus, hypothalamus [9,10], and adrenal glands, where it can directly interfere with the central and peripheral regulation of this major stress axis [6,11]. With advancing age, the HPA axis undergoes significant dysregulation marked by elevated basal cortisol, impaired negative feedback, altered glucocorticoid (GC) sensitivity, and altered stress responsiveness [12].

At the same time, reports have emerged regarding aging, the strongest risk factor for PD [1,13,14], as well as age-dependent impairment of the phagocytic capacity of microglia and peripheral macrophages regarding α-syn and cellular debris [15]. In PD, dysregulation of the peripheral immune system — particularly monocytes and T cells — has been reported multiple times, and the occurrence of both pathologic α-syn and a dysregulated monocyte compartment can reinforce each other to induce excessive inflammatory responses to α-syn [16]. Therefore, the age-related changes in the HPA axis may synergize with PD pathology to exacerbate neuroendocrine dysfunction and disease progression [12,17].

GCs are the principal mediators linking HPA axis activity to immune function [18]. Notably, reduced GC sensitivity of peripheral immune cells has also been reported in major depressive disorder (MDD), a psychiatric disorder frequently comorbid with PD [19], in which impaired GC responsiveness is thought to contribute to chronic low-grade inflammation and HPA axis dysregulation [20]. Given the substantial overlap between PD and depression with respect to HPA axis dysfunction and immune alterations, an impaired peripheral immune cell GC sensitivity may likewise represent an important downstream consequence of PD-associated pathology. However, direct evidence in the context of PD and α-syn pathology remains lacking [6,8,21].

The aim of the current study was therefore to investigate splenic GC sensitivity in a well-characterized α-syn overexpression transgenic mouse model (V1S/SV2) [22] across different ages and to compare the findings with physiological aging in wild-type (WT) C57BL/6J mice. Using an *ex vivo* splenocyte-based GC sensitivity assay, combined with analyses of HPA axis-associated parameters, splenocyte glucocorticoid receptor (GR) expression, and splenic GC signaling markers, we aimed to determine whether α-syn pathology alters peripheral immune-cell GC sensitivity beyond age-related effects.

## METHODS

### Generation and housing of transgenic animals

Generation of the V1S/SV2 mouse model and detailed phenotypic characterization have been described previously [22]. Animals were housed at the Animal Research Center of Ulm University under standardized conditions. Mice were group-housed in open polycarbonate type II long cages under controlled temperature and humidity conditions with a 12 h light/dark cycle and ad libitum access to water and standard mouse diet. Cages were enriched with nesting material and polycarbonate shelters.

### Weight data

After euthanasia by cervical dislocation, the adrenal glands (right and left) were removed and stored in 1 mL Hank’s balanced salt solution (HBSS; Gibco, Cat. No. 14175-095) on ice until weight determination. For adrenal gland weight determination, the surrounding fat was cut off under a binocular microscope. The spleens were extracted, freed from fat, and weighed, followed by direct splenocyte isolation.

### Splenocyte isolation

Following spleen extraction, splenocytes were isolated as previously described [23]. Briefly, whole spleens were homogenized through a 70 µm nylon cell strainer (Sarstedt, Cat. No. 83.3945.070) using a 2 mL syringe plunger in 20 mL ice-cold Hanks’ Balanced Salt Solution (HBSS; Gibco, Cat. No. 14175-095) and collected in 50 mL tubes. Cell suspensions were centrifuged at 600 × g for 12 min at 4°C, and the supernatant was discarded. Red blood cells were lysed by incubation in 1 mL lysis buffer (155 mM NH_4_Cl, 10 mM KHCO_3_, 10 mM EDTA, pH 8.0) for 2 min. Lysis was stopped by addition of 15 mL cold HBSS supplemented with 10% (v/v) FBS.

Cells were centrifuged again at 600 × g for 10 min at 4°C. After removal of the supernatant, cell pellets were resuspended in 5 mL cold HBSS and filtered once more through a 70 µm cell strainer, followed by an additional centrifugation step at 600 × g for 10 min at 4°C. The resulting cell pellet was resuspended in 15 mL cold RPMI++ medium. Splenocytes were counted manually using Neubauer-improved counting chambers. Cell suspensions were subsequently adjusted to a concentration of 5 × 10^6^ cells/mL by an additional centrifugation step and resuspension in the appropriate volume.

Freshly isolated splenocytes were used for GC sensitivity assays. Remaining cells were stored at −80°C in RLT lysis buffer (Qiagen, Cat. No. 74106) for subsequent qPCR analysis or cryopreserved in 90% (v/v) fetal bovine serum (FBS; Bio&Sell, Cat. No. FBS.HP.0500) and 10% (v/v) DMSO for later flow cytometry analysis.

### Cell culture and media

Isolated mouse splenocytes were cultured in Roswell Park Memorial Institute 1640 Medium (RPMI, Gibco, 21875-034) supplemented with 10% (v/v) fetal bovine serum (FBS, Bio&Sell, Cat. No. FBS.HP.0500) and 50U/ml Penicillin + 50µg/ml Streptomycin (hence RPMI(++)). Cells were cultivated in a 5% CO_2_ humidified incubator at 37 °C.

### GC sensitivity assay – splenocytes

GC sensitivity assays were performed using splenocytes isolated from V1S/SV2 mice at 6, 16, and 20 months of age, as previously described with modifications [23]. Lipopolysaccharide (LPS; Sigma-Aldrich, Cat. No. 27840) was diluted in RPMI++ medium and added to each well to yield a final concentration of 1 µg/mL. CORT (Sigma-Aldrich, Cat. No. C2505) stock solution was prepared by dissolving CORT powder in a mixture of 95% ethanol and RPMI++ medium to a concentration of 25 mM, and further diluted in RPMI++ medium to yield final well concentrations of 0, 0.05, 0.1, 0.5, and 5 µM.

For each condition, 45 µL RPMI++ medium, either without stimulation (basal) or supplemented with LPS, was added to wells of a white flat-bottom 96-well plate (Thermo Scientific, Cat. No. 136101), followed by 5 µL of the respective CORT dilution and 50 µL splenocyte suspension. For each mouse, 10 wells were prepared, comprising 5 basal wells and 5 LPS-stimulated wells. The final cell concentration in each well was 2.5 × 10^6^ cells/mL.

Cells were incubated for 24 h at 37°C and 5% CO_2_ in a humidified incubator. Cell viability was subsequently assessed using the CellTiter-Glo Luminescent Cell Viability Assay (Promega, Cat. No. G7571) according to the manufacturer’s instructions. Briefly, CellTiter-Glo substrate was added to each well, followed by incubation for 12 min at room temperature. Luminescence was measured as relative luminescence units (RLU) using a Victor X Light Luminescence Plate Reader (PerkinElmer).

For data analysis, luminescence values (RLU) obtained from unstimulated wells were subtracted from the corresponding LPS-stimulated wells (delta cell viability) for each treatment condition. The resulting delta RLU was set to 100% for the 0 µM CORT condition. A significantly lower cell viability of the 0.1 µM CORT condition compared to the CORT = 0 µM condition (set to 100%) was considered a functional indication of glucocorticoid sensitivity.

### Quantitative real-time PCR (qPCR) of *Fkbp5* and *Fkbp4* in murine splenocytes

Splenocytes were isolated from control and transgenic V1S/SV2 mice (n = 5 animals per group). Total RNA was extracted using the RNeasy Mini Kit (Qiagen, Cat. No. 74105; Qiagen, Hilden, Germany) according to the manufacturer’s instructions, including DNase treatment to remove genomic DNA contamination. RNA quantity and purity were assessed using a NanoDrop spectrophotometer.

cDNA was synthesized from 100 ng total RNA using the Bio-Rad iScript cDNA Synthesis Kit (Bio-Rad, Cat. No. 1708890) according to the manufacturer’s instructions. Quantitative real-time PCR (qPCR) was performed using iQ SYBR Green Supermix (Bio-Rad, Cat. No. 1708880) on a Bio-Rad CFX96 Real-Time PCR Detection System. Reactions were performed in a total volume of 20 µL and run in duplicate.

The following primer sequences were used: *Fkbp5* forward 5′-TGTTCAAGAAGTTCGCAGAGC-3′ and reverse 5′-CCTTCTTGCTCCCAGCTTT-3′; *Fkbp4* forward 5′-GACCGAGTCTTTGTCCACTACAC-3′ and reverse 5′-ATCCCAAGCCTTGATGACCTCC-3′.

PCR cycling conditions consisted of an initial denaturation step at 98°C for 30 s, followed by 35 amplification cycles with denaturation at 92°C for 30 s and annealing for 30 s using gene-specific annealing temperatures. Annealing temperatures were 64°C for *Fkbp5*, 69°C for *Fkbp4*, 69.1°C for *Gapdh*, and 72°C for *Ywhaz*. Melt curve analysis was performed from 65°C to 95°C to confirm amplification specificity.

Gene expression levels were normalized to the housekeeping genes *Gapdh* and *Ywhaz*. Relative gene expression was determined using the ΔΔCt method. Standard curves were generated to assess amplification efficiency. Data were analyzed using the Bio-Rad CFX Manager software (version 2.1.1022.0523).

### Western blot analysis

Protein samples were separated on SDS-polyacrylamide gels (5% stacking, 12% resolving) in a chamber filled with running buffer (25 mM Tris, 192 mM glycine, 1 g/L SDS). To enable comparison across membranes, a sample from the same control mouse was loaded on every gel as an inter-blot reference. Electrophoresis was run in two stages: 20 min at 80 V, followed by approximately 3 h at 100 V. Separated proteins were subsequently transferred onto nitrocellulose membranes (0.2 µm, Amersham Protran, 10600011) for 1.5 h at 25 V using the XCell II Blot Module (Invitrogen, EI0002) and transfer buffer (25 mM Tris, 192 mM glycine, 20% methanol). Membranes were blocked for 1 h at room temperature in 5% non-fat dry milk dissolved in TBS-T (1× TBS containing 0.05% Tween-20; Sigma-Aldrich, P9416) and incubated overnight at 4 °C with primary antibodies diluted in TBS-T (see below). After three washes in TBS-T, membranes were probed with the corresponding secondary antibodies (see below) diluted in 5% milk/TBS-T for 2 h at room temperature. Signals were developed with HRP substrate (SuperSignal West Pico PLUS, Thermo Fisher Scientific) and captured on a Fusion SoloS Imager (Vilber). Densiometric analysis was conducted with the Fusion software, and target protein signals were normalized to GAPDH and to the inter-blot reference sample to account for membrane-to-membrane variability.

Primary antibodies: Polyclonal rabbit anti-FKBP51 antibody (proteintech, 14155-1-AP; 1:500), rabbit anti-GAPDH (proteintech, 10494-1-AP; 1:10,000)

Secondary Antibodies: anti-rabbit IgG HRP (Promega, W401B, 1:10,000), anti-mouse IgG HRP (Promega, W402B, 1:10,000)

### Fecal CORT metabolite enzyme immunoassay

Fecal samples were collected during Rotarod behavioral testing of 5 mice expressing the transgene V1S/SV2 and 5 littermate control mice without transgene expression at 01:00 pm ±1, and processed as previously described [24]. Briefly, dry fecal pellets were homogenized thoroughly using a mortar and pestle. An aliquot of 0.05 g fecal material was extracted in 1mL of 80% methanol for 30 min on a shaking mixing block (BIOER Technology, MB-101). Samples were subsequently centrifuged at 2500 x g for 15 min. Following centrifugation, 100 µL aliquots of the supernatant were collected in duplicate and stored at −20°C until fecal CORT metabolite measurement using an established 5α-pregnane-3β,11β,21-triol-20-one enzyme immunoassay (EIA). This assay has been established and extensively validated for monitoring glucocorticoid metabolites in fecal samples of mice [24,25].

### Flow cytometry

Frozen murine splenocytes of 5 transgenic V1S/SV2 and 5 control mice in 90% FBS and 10% DMSO were rapidly thawed in a 37°C water bath and immediately transferred into pre-warmed RPMI++. Cells were centrifuged at 400 x g for 5–10 minutes to remove residual DMSO, and the resulting pellet was resuspended for subsequent staining. For immunophenotyping of different cell populations, 1-2 × 10^6^ cells were washed with PBS and stained with ViaDye Red (1:10,000 dilution) for 15 minutes at room temperature in the dark to assess viability. Following a wash with FACS buffer (PBS + 0.5% BSA), Fc receptors were blocked using FcR blocking reagent (Miltenyi Biotec) for 10 minutes at 2-8°C. Surface antigens were then labeled by incubating cells with an antibody master mix for 30 minutes at 2-8°C. For the detection of intracellular targets, including Glucocorticoid Receptor (GR) and FoxP3, cells were fixed and permeabilized using the Transcription Factor Staining Buffer Set (Miltenyi Biotec) according to the manufacturer’s instructions. Intracellular staining was performed in permeabilization buffer for 30 minutes at 2-8°C in the dark. After a final wash, cells were resuspended in 300 µl of FACS buffer. Data were acquired on a four-laser Cytek Aurora spectral flow cytometer (Cytek Biosciences).

### Antibodies used

CD45-Alexa Fluor 700 (103128, Biolegend, 1:400), CD19-BV650 (115541, Biolegend, 1:200), CD3-V500 (560771, BD Biosciences, 1:200), CD4-PerCP (553052, BD Biosciences, 1:200), CD8-BV570 (100740, Biolegend, 1:100), CD11b-eFluor450 (40011292, Invitrogen, 1:200), CD11c-APC (550261, BD Biosciences, 1:200), Ly6C-BV605 (128036, Biolegend, 1:100), Ly-6G-Spark UV387 (127678, Biolegend, 1:50), MHC-II-BUV496 (750281, BD Biosciences, 1:200), F4/80-BV421 (123132, Biolegend, 1:30), CD206-BV785 (141729, Biolegend, 1:100), Glucocorticoid receptor (GR)-Alexa Fluor 488 (53-6189-82, Invitrogen, 1:100), Mouse IgG2a, kappa isotype control for GR -AF488 (53-4724-80, Invitrogen, 1:100), Live-dead ViaDye Red (R7-60008, Cytek Bioscience, 1:10,000), CD3-eFluor 450 (48-0031-82, Invitrogen, 1:100), CD4-PerCP (553052, BD Biosciences, 1:200), CD25-PE (12-0251-81B, Invitrogen, 1:100), FoxP3-APC (17-5773-82, Invitrogen, 1:40)

### Software and data analysis

Wet-lab statistical analyses and data visualization were performed using GraphPad Prism (version 11.0.0; GraphPad Software). Flow cytometry data were analyzed using FlowJo (version 11.1.1; BD Biosciences). Prior to hypothesis testing, the normality of data distribution was assessed using the Kolmogorov–Smirnov test with Lilliefors’ correction. Potential outliers within normally distributed datasets were identified by Grubbs’ test and excluded from further analysis. For comparisons between two independent groups with normally distributed data, the unpaired two-tailed Student’s t-test was applied when the F-test confirmed equal variances, or the unpaired t-test with Welch’s correction when variances were significantly unequal; for paired comparisons within the same animals, the paired t-test was used. For all two-group and three-group comparisons of normalized dose–response data across the full CORT concentration range, ordinary two-way ANOVA followed by Tukey’s multiple comparisons test was performed. When the assumption of normality was not met in single-concentration comparisons, the Mann–Whitney U test was applied for two independent samples and the Wilcoxon signed-rank test for two paired samples. All data are presented as mean ± SD with individual values shown as dots. The threshold for statistical significance was set at p ≤ 0.05.

### Large Language models (LLMs) and generative AI

AI-assisted writing tools (Claude, Anthropic; ChatGPT, OpenAI) were used for language editing and text formulation. All scientific content, data interpretation, and conclusions are solely the responsibility of the authors.

## RESULTS

### V1S/SV2 mice develop splenic GC resistance at 16 months

To characterize the effect of aging on splenic *ex vivo* GC resistance, we assessed splenocyte sensitivity to CORT in an aging cohort of WT mice at 6, 16, and 20 months of age. LPS-induced increase in proliferation and viability, and the counteracting effect of CORT in this system, was used as a proxy for GC sensitivity. In this cohort, LPS robustly increased splenocyte viability compared to the basal condition at all ages (Fig. 1a, left; *p = 0.02, **p = 0.008, **p = 0.0039, Wilcoxon test), confirming the overall functionality of the assay. The difference between LPS-stimulated and basal conditions (delta cell viability) did not significantly vary with age, indicating a comparable LPS response across groups (Fig. 1a, middle; p > 0.05, Mann-Whitney test). To evaluate GC responsiveness, the delta cell viability was normalized to the 0 µM-CORT condition across the full CORT dose range. Our data revealed an age-associated reduction in GC sensitivity: two-way ANOVA identified significant main effects of both age (***p = 0.0004) and CORT treatment (***p = 0.0001), with splenocytes from 20-month-old mice suppressed more weakly by CORT than those from 6-month-old mice at 0.1 and 0.5 µM CORT (Fig. 1a, right; adjusted *p = 0.011 and adjusted **p = 0.007, Tukey’s multiple comparisons). Together, these findings indicate a progressive decline in GC sensitivity of splenocytes with age in WT mice.

**Fig. 1:**
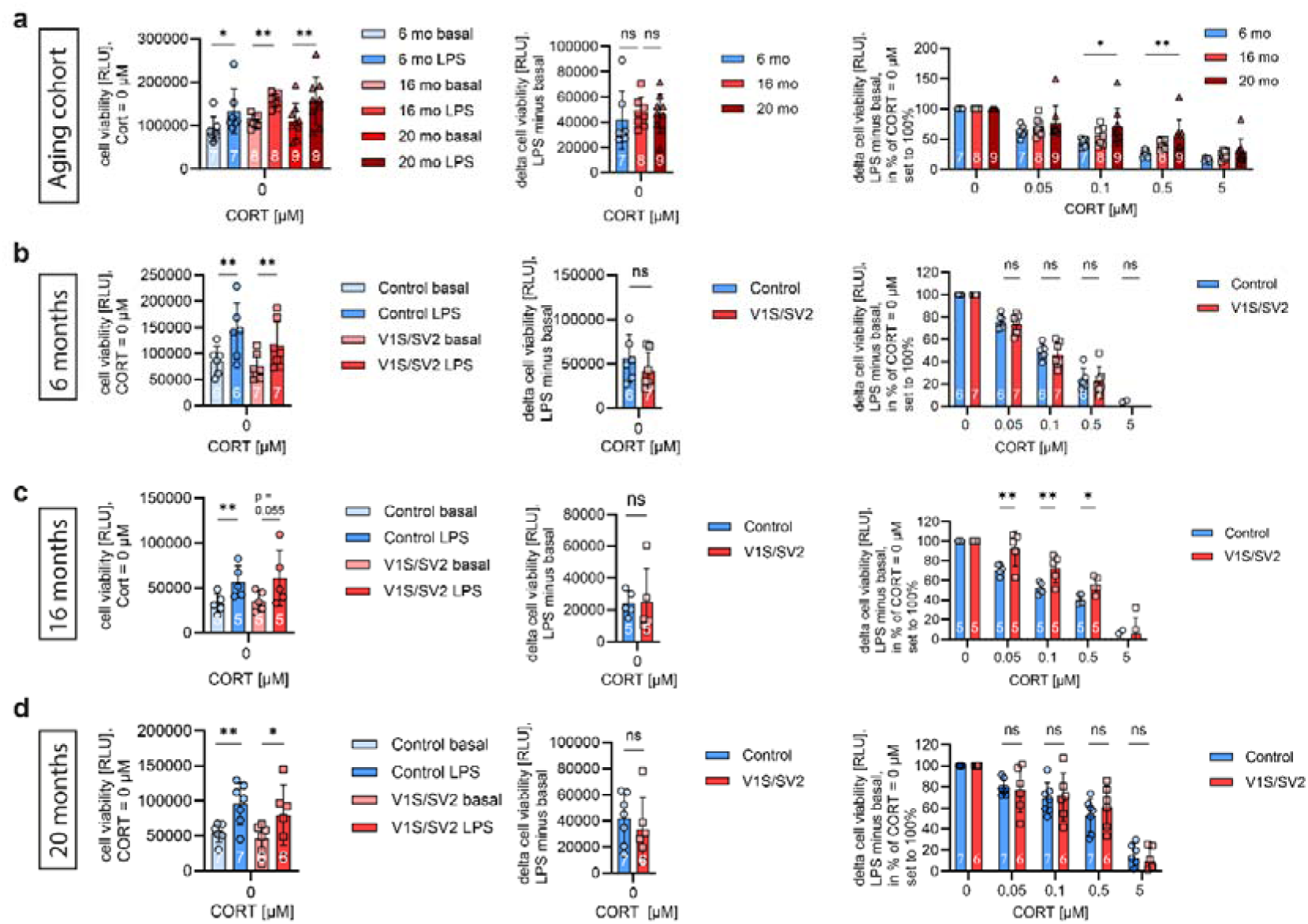
Dysregulated splenic glucocorticoid (GC) sensitivity in aging and the V1S/SV2 Parkinson’s disease (PD) mouse model at 16 months. GC sensitivity was assessed in freshly isolated splenocytes from a separate wild-type (WT) aging cohort (a) or V1S/SV2 mice and littermate controls across three age groups (b-d). Splenocytes were stimulated with lipopolysaccharide (LPS; 1 µg/mL) and cultured for 24 h with increasing concentrations of corticosterone (CORT; 0, 0.05, 0.1, 0.5, and 5 µM). Cell viability was quantified using the CellTiter-Glo luminescent assay (relative luminescence units, RLU). (a), WT aging cohort (6, 16, and 20 months). Left: Absolute cell viability (RLU) under unstimulated (basal) and LPS-stimulated conditions at CORT = 0 µM. Middle: Delta cell viability (LPS minus basal, RLU) at CORT = 0 µM across age groups. Right: Delta cell viability normalized to the condition without CORT (CORT = 0 µM, set to 100%) across the full CORT dose range. (b), 6-month cohort, (c), 16-month cohort, and (d), 24-month cohort (Control vs. V1S/SV2). For each cohort: Left: Absolute cell viability (RLU) at CORT = 0 µM for basal and LPS conditions. Middle: Delta cell viability (LPS minus basal, RLU) at CORT = 0 µM. Right: Normalized delta cell viability to CORT = 0 µM (set to 100%) across the dose range. Data are shown as mean ± SD with individual data points, *p<0.05, **p<0.01, ***p<0.001; n = 5 – 9

Given this age-dependent decline, we next examined GC signaling in the V1S/SV2 mouse model to determine whether genotype-specific effects extend beyond aging. LPS consistently increased cell viability relative to the basal condition in both V1S/SV2 mice and littermate controls at 6 months (**p = 0.004 and **p = 0.002, respectively), at 16 months (**p = 0.002 and p = 0.056, respectively), and at 20 months (**p = 0.001 and *p = 0.019, respectively; Fig. 1b–d left, paired t-test). The difference between LPS-stimulated and basal conditions (delta cell viability) did not significantly vary in V1S/SV2 compared to controls in all age groups, indicating a comparable LPS response across groups (Fig. 1b-d middle; p > 0.05, unpaired t test). To evaluate GC responsiveness, the delta cell viability was normalized to the 0 µM-CORT condition across the full CORT dose range. At 6 months, V1S/SV2 splenocytes did not differ from controls across the entire CORT dose range (Fig. 1b right; adjusted p > 0.05, two-way ANOVA with Tukey’s multiple comparisons test). A significant main effect of CORT treatment emerged (***p < 0.0001), whereas genotype had no significant effect. In contrast, at 16 months, V1S/SV2 splenocytes developed a pronounced GC resistance. A two-way ANOVA followed by Tukey’s multiple comparisons test identified significant main effects of genotype and CORT treatment (both ***p < 0.0001), with higher normalized delta cell viability in V1S/SV2 mice than controls at 0.05 µM (Fig. 1c right; adjusted **p = 0.002), 0.1 µM (adjusted **p = 0.0035), and 0.5 µM CORT (adjusted *p = 0.026). By 20 months, this difference was no longer detectable, as both genotypes displayed a similarly attenuated CORT response (Fig. 1d right; adjusted p > 0.05, two-way ANOVA with Tukey’s multiple comparisons test). Two-way ANOVA revealed a significant main effect of CORT treatment (***p < 0.0001) but no significant effect of genotype. In other words, by 20 months, control mice had aged into the same GC-resistant state already exhibited by V1S/SV2 mice at 16 months, masking the genotype effect. The 16-month time point, therefore, defined a critical window in which genotype-driven alterations in GC sensitivity are detectable.

### GC resistance in 16-month-old splenocytes occurs despite intact HPA axis output and stable FKBP51 and GR protein levels

To investigate the mechanisms underlying the splenic GC resistance phenotype observed in 16-months V1S/SV2 mice, we examined key components of HPA-axis function and peripheral GC signaling, including FKBP51 protein abundance as a cochaperone of GR, splenic gene expression, and GR protein abundance at this time point (Fig. 2). Relative spleen weight (Fig. 2a), relative adrenal-gland weight (Fig. 2b), and fecal CORT concentration (Fig. 2c) did not differ between V1S/SV2 mice and controls (p > 0.05, unpaired t-test). Lymphoid-organ size and adrenal output were therefore preserved, and the immune cell GC resistance phenotype could not be attributed to chronic hyper- or hypocortisolism.

**Fig. 2:**
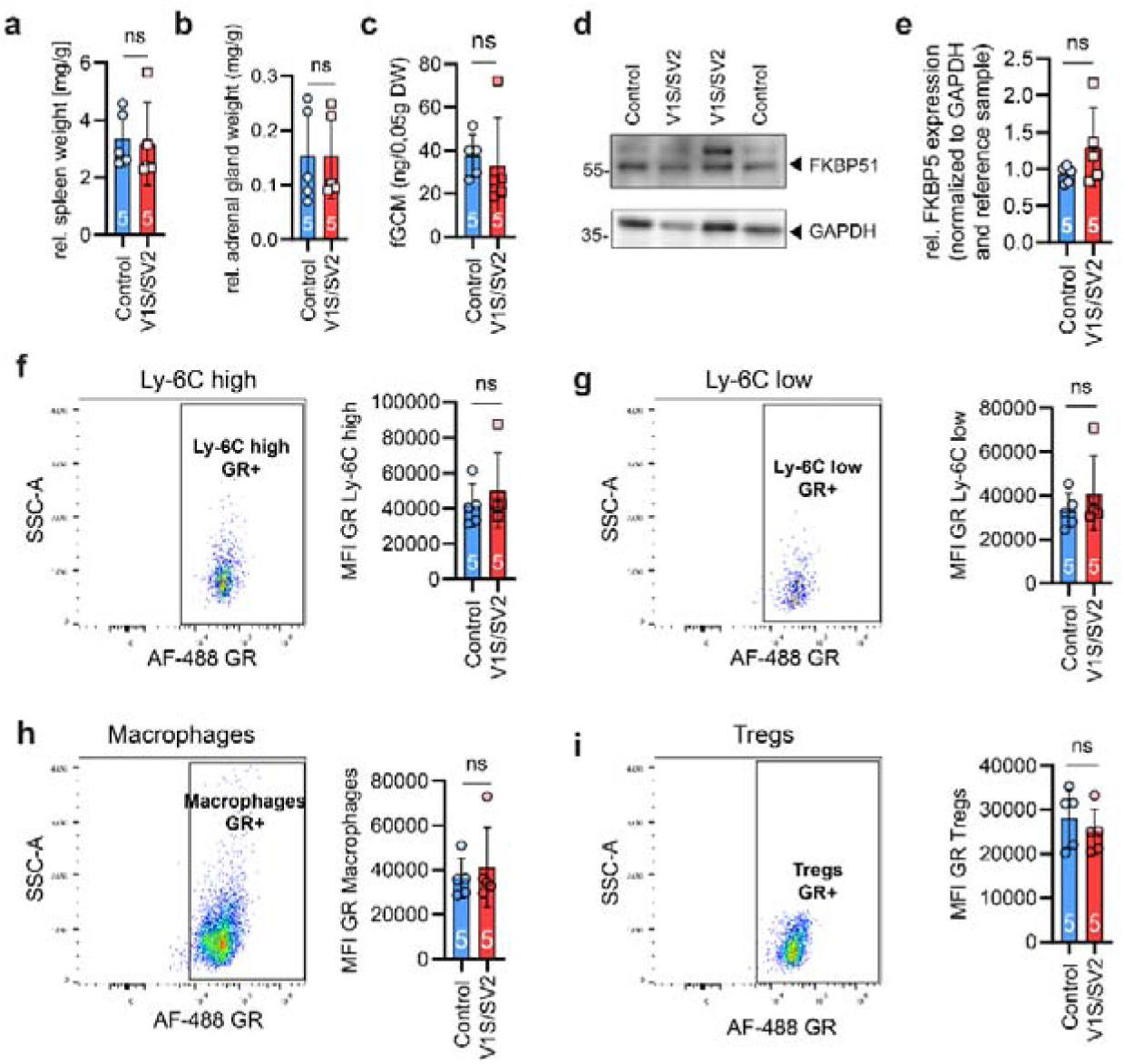
Splenic glucocorticoid (GC) resistance in 16-month-old V1S/SV2 mice is not explained by changes in hypothalamic–pituitary–adrenal (HPA)-axis output, FKBP51 protein abundance, or glucocorticoid receptor (GR) protein expression. Characterization of HPA axis and glucocorticoid signaling markers in 16-month-old V1S/SV2 mice and wild-type (WT) littermate controls — the only age group showing a splenic GC resistance phenotype (Fig. 1c). (a), Relative spleen weight (mg/g body weight) for V1S/SV2 mice and the WT control group after 16 months. (b), Relative adrenal gland weight (mg/g body weight) for V1S/SV2 mice and the WT control group after 16 months. (c), Fecal corticosterone metabolite concentration (fCORT; ng per 0.05 g dry weight, feces collected at 01:00 pm ±1, measured by 5α-pregnane-3β,11β,21-triol-20-one enzyme immunoassay [EIA]) for V1S/SV2 mice and the WT control group after 16 months. (d), Relative FKBP51 expression normalized to GAPDH and the reference sample measured by Western blot for V1S/SV2 mice and the WT control group after 16 months. (e-h), Glucocorticoid receptor (GR) protein expression in splenic immune cell subsets: Ly-6C^high^ monocytes (e), Ly-6C^low^ monocytes (f), Macrophages (CD11b+ F4/80+) (g), Regulatory T cells (CD4+ CD25+ FoxP3+; Tregs) (h), measured by flow cytometry. For each subset, a representative pseudocolor flow cytometry plot (GR-AF488 on the x-axis) is shown on the left and the quantification of the GR median fluorescence intensity (MFI) on the right. Data are presented as mean ± SD with individual data points for each animal, *p<0.05, **p<0.01, ***p<0.001, n = 5 control, 5 V1S/SV2.

To assess downstream GR signaling activity, we quantified splenic FKBP51 protein by Western blot. FKBP51 — a robust GR target and a sensitive readout of GR transcriptional activity [26] — did not differ significantly between V1S/SV2 mice and controls at 16 months (Fig. 2d; p > 0.05, unpaired t-test with Welch’s correction). At the mRNA level, however, splenic *Fkbp5* mRNA was significantly reduced in V1S/SV2 splenocytes at this time point (Fig. S1b; *p = 0.03, Mann-Whitney test). The related GR co-chaperone *Fkbp4* showed no difference compared to controls at the same time point (Fig. S1b; p > 0.05, Mann-Whitney test). The *Fkbp5* effect was specific to 16 months: at 6 and 20 months, *Fkbp4* and *Fkbp5* levels were comparable between V1S/SV2 and control splenocytes, both in disease cohorts and in the WT aging cohort (Fig. S1a, c, d; p > 0.05, unpaired t test).

To determine whether the GC resistance phenotype was accompanied by changes in GR protein expression, we quantified GR by intracellular flow cytometry in the major splenic immune subsets at 16 months. GR median fluorescence intensity (MFI) did not differ between V1S/SV2 mice and controls in Ly-6C^high^ monocytes, Ly-6C^low^ monocytes, F4/80^+^ macrophages, or CD4^+^ CD25^+^ FoxP3^+^ regulatory T cells (Fig. 2e– h; p > 0.05, Mann-Whitney test). Extended phenotyping with the same panel showed unchanged GR levels in CD4^+^ T cells, dendritic cells (CD11c^+^ MHC-II^+^), and neutrophils (Ly-6C^+^ Ly-6G^+^) (Fig. S2a–c; gating strategy and FMO controls in Fig. S2d–I; p > 0.05, Mann-Whitney test).

Taken together, V1S/SV2 mice developed splenic GC resistance at 16 months, accompanied by a selective reduction in *Fkbp5* mRNA expression, and occurring independently of systemic HPA axis alterations, FKBP51 protein changes, or GR downregulation in splenic immune subsets.

## DISCUSSION

Despite substantial evidence for systemic immune alterations [8,21], GC sensitivity of peripheral immune cells in PD has not been investigated. We therefore assessed *ex vivo* GC sensitivity of isolated and LPS-stimulated splenocytes [23] in the previously described V1S/SV2 α-syn transgenic mouse model [22] across aging. V1S/SV2 mice developed a transient GC resistance phenotype at 16, but not 6 and 20, months of age, characterized by a reduced CORT-mediated suppression of delta (i.e., LPS – basal) splenocyte viability despite unaffected adrenal output as well as GR and FKBP51 protein levels. Importantly, as aging alone progressively reduced *ex vivo* GC sensitivity of isolated and LPS-stimulated splenocytes in WT mice, the lack of *ex vivo* splenic GC resistance in 20-month-old V1S/SV2 mice might be due to a general age-related decline in GC sensitivity masking possible genotype effects.

The emergence of splenic GC resistance at 16 months occurs during a critical phase of α-syn-driven disease progression in V1S/SV2 mice. This time point corresponds to a stage of progressive α-syn accumulation, including established oligomeric pathology in multiple forebrain and midbrain regions [22]. In the closely related S1/S2 model, a substantial transcriptomic overlap with α-syn pathology-associated genes was observed [13]. At this stage, V1S/SV2 mice additionally exhibit motor and olfactory impairments that are absent at 6 months of age [13,22]. Together, these findings identify the 16-month time point as a stage of biologically meaningful α-syn-driven circuit dysfunction that coincides with a functionally measurable alteration in ex vivo GC responsiveness of peripheral immune cells. Peripheral immune dysfunction is a hallmark of PD [21,27]. Our findings extend these observations by demonstrating that α-syn pathology is associated not only with immune alterations at the cellular level, but also with impaired responsiveness to anti-inflammatory GCs, the principal endocrine mediators linking HPA axis activity to immune function [18].

Peripheral immune GC resistance may represent both a consequence of α-syn-driven pathology and a mechanism that further amplifies neuroinflammatory disease processes in PD. The *ex vivo* GC resistance observed in peripheral immune cells from V1S/SV2 mice is consistent with impaired *ex vivo* GC responsiveness described in multiple conditions associated with chronic stress, HPA axis dysregulation, and systemic immune activation. For instance, reduced *ex vivo* GC sensitivity of peripheral immune cells has been described in patients diagnosed with MDD [20], post-traumatic stress disorder [28], and in rodent models of psychosocial stress, including the social disruption (SDR) paradigm and the chronic subordinate colony housing (CSC) paradigm [23,29,30]. This context is particularly relevant because approximately 40% of PD patients suffer from comorbid depression, which frequently emerges during the prodromal phase, suggesting overlapping upstream mechanisms [6]. We propose that by failing to suppress α-syn -driven immune activation, GC-resistant immune cells sustain a pro-inflammatory state that promotes microglial activation, oxidative stress, mitochondrial dysfunction, and ultimately contributes to neurodegenerative processes seen in depression and/or PD.

The fact that splenic *ex vivo* GC resistance was first detected in V1S/SV2 mice at an age of 16 months, and not in WT mice at an age of 20 months, suggests that α-syn pathology accelerates an aging-associated decline in splenic GC responsiveness. A general age-dependent decline in GC sensitivity of peripheral immune cells is consistent with earlier studies in humans [12,17]. Moreover, as isolated splenocytes from both V1S/SV2 mice and controls at the age of 20 months showed a comparably low splenic *ex vivo* GC sensitivity, it is likely that a general age-related decline in GC sensitivity masked possible genotype effects; instead, isolated splenocytes from V1S/SV2 mice at the age of 20 months have overcome GC resistance seen at the age of 16 months. As aging represents the strongest risk factor for PD [13,14] and independently promotes GC resistance of peripheral immune cells [17], both processes likely converge at advanced age. Thus, splenic GC resistance is not unique to α-syn pathology; rather, α-syn pathology appears to accelerate its onset relative to physiological aging. Together, the WT and transgenic aging data support a model in which early α-syn pathology accelerates an age-associated decline in peripheral GC responsiveness, with the 16-month time point representing a critical window for PD progression and GC resistance.

The observed GC resistance phenotype at 16 months appears to arise independently of systemic HPA axis alterations and GR dysregulations on immune cells. Unaffected relative spleen and adrenal gland weights as well as unaltered fecal CORT concentrations indicate that systemic HPA axis output was intact — excluding chronic hypercortisolism-driven GR downregulation as a contributing factor [8,18]. FKBP51 protein — a sensitive bidirectional readout of GR transcriptional activity that is both a GR target gene and a negative regulator of GR ligand-binding affinity [26] — was likewise unchanged. Intracellular GR quantification across Ly-6C^high^ monocytes, Ly-6C^low^ monocytes, F4/80^+^ macrophages, and CD4^+^CD25^+^FoxP3^+^ Tregs further showed no difference in GR median fluorescence intensity, ruling out receptor-level downregulation. Together, these findings suggest that the observed splenic *ex vivo* GC resistance arises downstream of receptor expression, with impaired GR nuclear translocation representing a plausible mechanism. In the SDR model of psychosocial stress, splenic GC resistance developed despite intact GR protein expression; the molecular mechanism was localized to CD11b^+^ cells and consisted of defective GR nuclear translocation following GC stimulation, resulting in impaired suppression of NF-κB-driven pro-inflammatory gene transcription as a consequence of LPS stimulation [30]. This mechanism is particularly relevant given our finding that GR expression is intact in CD11b^+^ cells and other splenic immune cell populations, suggesting that GC resistance in V1S/SV2 splenocytes may similarly operate at the level of GR activation and nuclear import rather than receptor abundance. A related mechanism has been described in the CSC model, where polymorphonuclear myeloid-derived suppressor cells (PMN-MDSCs) were identified as the cellular driver of GC resistance through a TLR4/NF-κB -dependent pathway [29]. The latter is potentially relevant, given the known dysregulation of myeloid populations under conditions of α-syn -driven neuroinflammation [27].

Our data integrate into a broader framework linking α-syn pathology, HPA axis dysfunction, and impaired GC signaling in PD. Plasma cortisol is consistently elevated in PD patients, appears independent of dopaminergic treatment and disease duration, and has been proposed to reflect a fundamental disruption of HPA axis homeostasis [8,11]. Lewy body pathology further suggests that α-syn directly affects HPA-axis-associated structures [9,11]. In a BAC transgenic α-syn rat model, chronic CORT administration aggravated phospho-Ser129 α-syn accumulation in the hypothalamus and hippocampus, while α-syn overexpression itself reduced hypothalamic CRF expression and altered baseline HPA-axis parameters, highlighting the bidirectional interaction between HPA axis dysfunction, altered GC signaling, and α-syn pathology [31]. Reduced GR expression in the substantia nigra of postmortem PD brains [6] and the neuroprotective role of microglial GR in rodent PD models [32] indicate that GC signaling competence is broadly compromised in PD across both the CNS and the peripheral immune system.

Targeting dysfunctional GR signaling has recently emerged as a promising therapeutic concept across multiple neurodegenerative disorders. In PD, dexamethasone exerted neuroprotective effects in a neuromelanin-based model by suppressing neuroinflammation [33]. Related findings from Amyotrophic lateral sclerosis [34] and Huntington’s disease, where GR antagonism attenuates neurodegeneration [35], further support GR signaling competence as a therapeutically relevant target across the neurodegenerative spectrum.

In summary, our data identify a disease-specific window of splenic *ex vivo* GC resistance in V1S/SV2 mice that precedes the equivalent age-related decline in WT controls and arises independently of systemic HPA axis output. These findings position peripheral immune GC resistance as a candidate mechanism by which α-syn pathology may amplify neuroinflammatory disease progression in PD.

## ACKNOWLEDGEMENTS

We would like to thank R. Bück, P. Hornischer, and U. Binder for their excellent technical support, as well as the ULMTeC Core Facility Immune Monitoring of the Medical Faculty at Ulm University for providing support and instrumentation funded by the Deutsche Forschungsgemeinschaft (DFG, German Research Foundation) - Project number 514808265.

## Statement of Ethics

Mouse experiments were performed in accordance with the German Animal Welfare Act (Tierschutzgesetz) and in line with the local guidelines of the Animal Research Center, Ulm University. This study was approved by the Federal Animal Care and Use Committee (Regierungspräsidium Tübingen, Germany), approval numbers o.190-12.TschB:W, 1576.TschB:S/C/H and o.165-5.TschB:C. All efforts were made to minimize the number of animals used and their suffering.

## Author Contributions

Project conception and design were carried out by DR, VB, SOR, and KMD. Experiments were performed, and data were analyzed by DR, with mouse experiments supported and conducted by VB. DL, VG, and JK contributed intellectual input. CT provided intellectual input and assisted with fecal CORT metabolite measurements. The manuscript was written by DR, SOR, and KMD and critically reviewed and approved by all authors.

## Data Availability Statement

All data generated or analyzed during this study are included in this article and its supplementary material source data files. Further enquiries can be directed to the corresponding author.

