## Supplementary information for "Dysregulated splenic glucocorticoid sensitivity in aging and an α-synuclein transgenic mouse model of Parkinson’s disease"

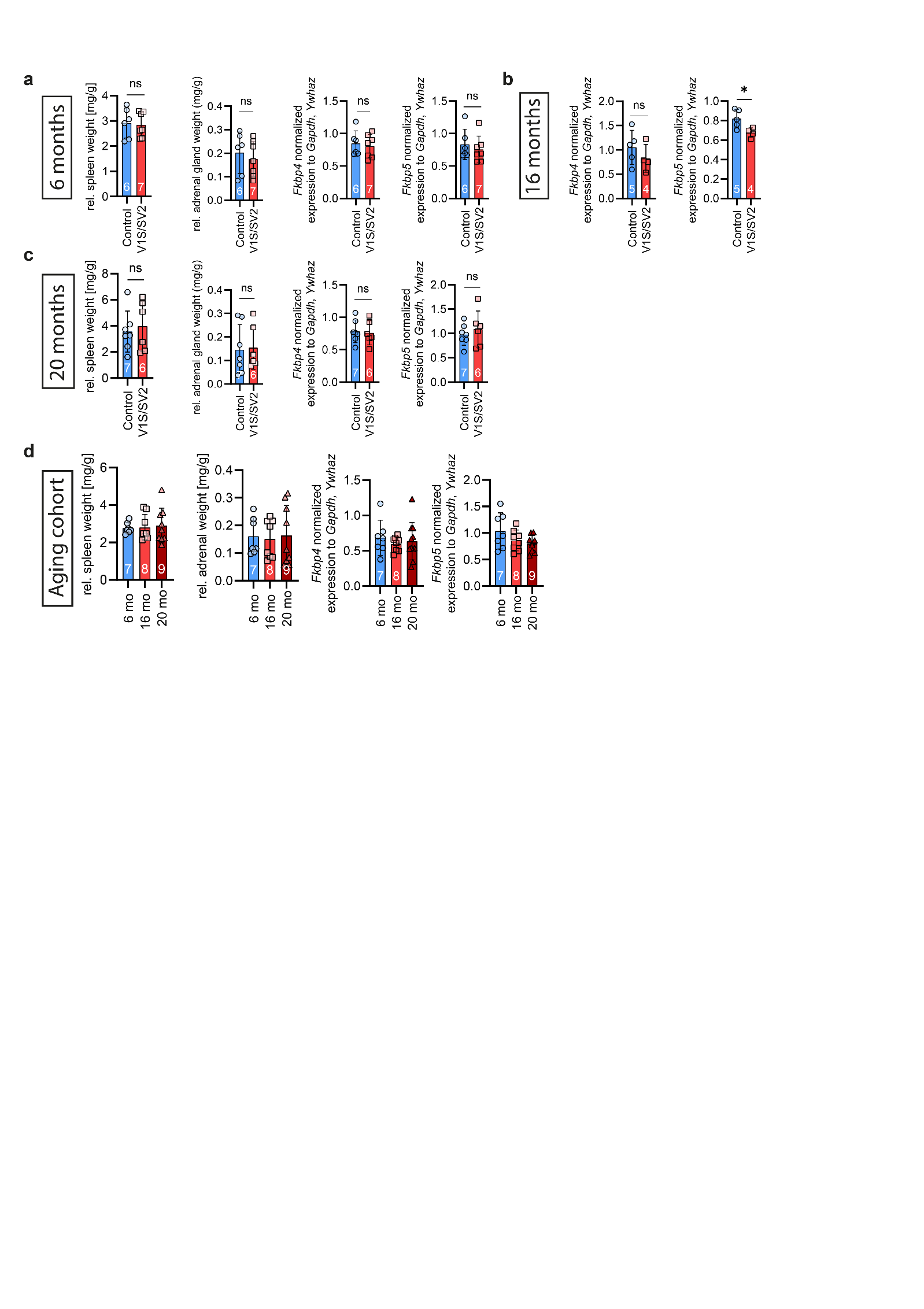

**Figure S1: Spleen and adrenal gland weights and *Fkbp4* / *Fkbp5* expression across ages in V1S/SV2 and wild-type (WT) mice. (a)**, 6-month cohort (n = 6 controls vs. 7 V1S/SV2). From left to right: relative spleen weight (mg/g), relative adrenal gland weight (mg/g), Relative splenic *Fkbp4* and *Fkbp5* mRNA expression (normalized to housekeeping genes *Gapdh* and *Ywhaz*) assessed by RT-qPCR. (**b)**, 16-month cohort (n = 5 controls vs. 4 V1S/SV2). From left to right: Shown is the relative *Fkbp4* and *Fkbp5* mRNA expression (normalized to housekeeping genes *Gapdh* and *Ywhaz*) assessed by RT-qPCR. (**c)**, 20-month cohort (n = 7 controls vs. 6 V1S/SV2). Same panel layout and statistics as in (a). (**d),** WT aging cohort with n = 7-9 animals per group (6, 16, and 20 months). Data are presented as mean ± SD with individual data points for each animal, *p<0.05, **p<0.01, ***p<0.001.

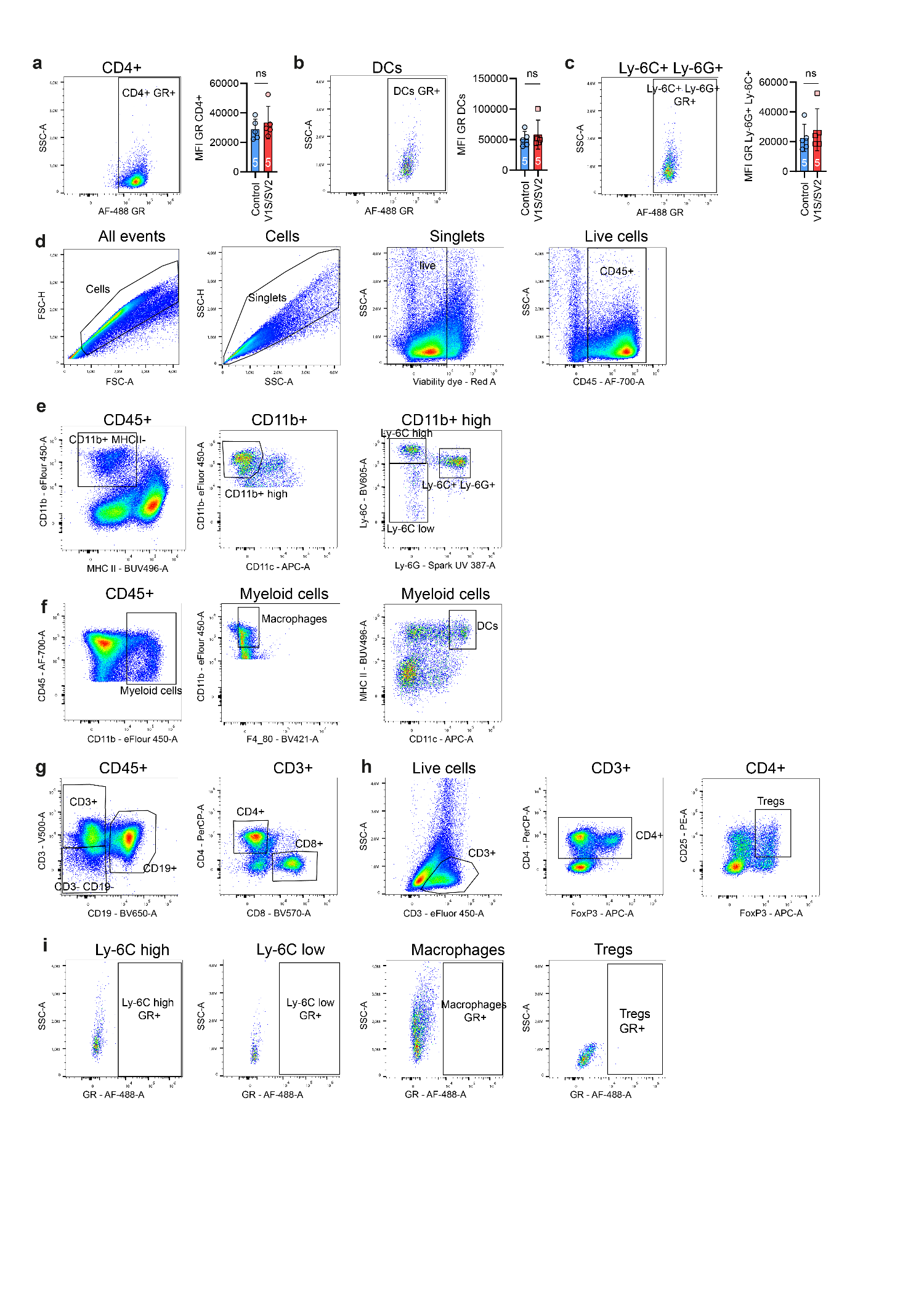

**Figure S2: Extended flow cytometric characterization of splenic glucocorticoid receptor expression and gating strategy.**

Flow cytometry analysis of splenocytes from 16-month-old V1S/SV2 mice and controls, extending the main-figure GR quantification (Fig. 2e–h) to additional immune cell subsets and documenting the full gating hierarchy and FMO controls. (**a-c)**, GR median fluorescence intensity (MFI) in additional splenic immune cell subsets. For each subset, a representative pseudocolor flow cytometry plot (GR-AF488 on x-axis) is shown on the left and the MFI quantification on the right. (a) CD4+ T cells. (b) Dendritic cells (DCs; CD11c+ MHC-II+). (c) Ly-6C+ Ly-6G+ neutrophils. Data are presented as mean ± SD with individual data points for each animal, p > 0.05, n = 5 controls vs. 5 V1S/SV2 at 16 months. **(d)**, General gating strategy on viable single cells and CD45+ cells. Sequential gating from All events → Cells (FSC-A vs. SSC-A) → Singlets (SSC-A vs. SSC-H) → Live cells (viability dye–negative) → CD45+ cells. **(e)**, Myeloid gating in the CD45+ compartment: CD11b+ → CD11b^high^ → Ly-6C^high^ / Ly-6C^low^ subsets. (**f)**, Further myeloid gating: CD45+ → myeloid cells → macrophages (F4/80+) and DCs (CD11c+ MHC-II+). **(g)**, T cell gating: CD45+ → CD3+ → CD4+/CD8+ cells. **(h)**, Regulatory T cell gating (Tregs): Live cells → CD3+ → CD4+ → CD25+ FoxP3+ (Tregs). **(i)**, Fluorescence-minus-one (FMO) controls for GR (AF488) staining in Ly-6C^high^, Ly-6C^low^, macrophages, and Tregs, used to define GR+ gate boundaries.
